# From Bad to Worse: Carryover Effects Alter Responses to Multiple Stressors

**DOI:** 10.64898/2026.10.09.757873

**Authors:** Charlotte Carrier-Belleau, Paul Brooks, Lauric Feugere, Philippe Archambault, Piero Calosi, Jennifer Coughlan, Alexina Trudelle, Tasman Crowe, Julien Gigault

## Abstract

Ecological responses to environmental stressors are increasingly shaped by interactions among anthropogenic and climatic drivers, yet the influence of prior exposure history on subsequent stress responses remains poorly integrated into multiple stressor ecology. Using the blue mussel (*Mytilus edulis*) as a model system, we experimentally tested whether prior exposure to environmental nanoplastics modifies responses to subsequent stressors, including copper contamination, nutrient enrichment, and salinity fluctuations, individually and in combination. Nanoplastic pre-exposure reduced survival across subsequent treatments and altered stressor interaction outcomes at the community level, shifting responses toward synergistic effects that amplified metabolic costs. These results demonstrate that exposure history can modify both the magnitude and direction of multi-stressor effects, with consequences extending beyond individual performance to ecosystem functioning. By incorporating temporal exposure dynamics into multiple stressor research, our study highlights how ecological responses to global change depend not only on the identity and intensity of stressors, but also on prior environmental conditions shaping subsequent organismal and ecosystem health status and responses.

**Open Research Statement:** The data and R scripts supporting the findings of this study will be deposited in a public repository upon acceptance

## 1. Introduction

Natural ecosystems are increasingly affected by rapid environmental changes driven by climate variability and human activities (Trégarot et al., 2024). Rather than occurring as isolated events, these changes expose organism to sequences of natural and anthropogenic stressors that vary in intensity, duration, and timing across space and time. These stressors can interact to create additive, synergistic (amplified), or antagonistic (attenuated) effects, often generating unexpected ecological outcomes that complicate conservation and management efforts (Côté et al., 2016). However, despite rapid advances in multiple stressor ecology, most experimental studies still assume that organisms encounter multiple stressors with identical environmental histories, an assumption that rarely reflects conditions experienced in nature (Jackson et al., 2021).

Temporal variation is increasingly recognized as a fundamental component of environmental change in the Anthropocene, where stressors are becoming more frequent, episodic, and recurrent (Ryo et al., 2019). Under these conditions, previous environmental experiences can influence subsequent performance through carryover effects, whereby the consequences of an earlier exposure persist after the original stressor has ceased (Harrison et al., 2011). Carryover effects can arise through persistent physiological changes, including oxidative stress, DNA damage, or disrupted energy balance, altering an organism’s capacity to respond to future environmental stress (Harrison et al., 2011). Consequently, environmental history may not only affect organismal performance but also modify the magnitude and even the nature of the interactive effects of multiple stressors. Yet, despite their potential importance, carryover effects remain largely overlooked in multiple stressor research, limiting our ability to predict ecological responses in natural ecosystems (Brooks & Crowe, 2019).

Coastal marine ecosystems provide an ideal model system in which to investigate these processes as organisms are routinely exposed to fluctuating environmental conditions and multiple anthropogenic stressors. Specifically, copper contamination, nutrient enrichment, and salinity fluctuations are widespread stressors that influence physiology, growth, reproduction, and survival across a broad range of aquatic taxa (Butt et al., 2022; Carrier-Belleau et al., 2024; Krishna et al., 2025). Beyond their effects on individual organism, these stressors can also alter the ecological functions performed by foundation species, with cascading consquences for community assumble and ecosystem functioning (Lauzon et al. 2026). Although these stressors affect organisms through different mechanisms, they often converge on shared physiological processes (e.g., energy metabolism, ion regulation, and oxidative balance), creating the potential for non-additive (interactive) effects. For example, reduced salinity can enhance copper toxicity (Grosell et al., 2007), while nutrient enrichment can modify organismal responses to fluctuating salinities (Carrier-Belleau et al., 2021). Whether these interactive effects are modified by previous environmental history, however, remains largely unknown.

Among emerging environmental stressors, nanoplastics (NP) are receiving increasing attention because of their widespread occurrence, high reactivity, and capacity to interact with other contaminants (Gigault et al., 2018, 2021; Cai et al. 2026). Experimental studies have demonstrated that NP can impair growth, reduce energy reserves, and increase mortality across a broad range of aquatic organisms (Blancho et al., 2021; Rochman et al., 2019; Trevisan et al. 2022). Their persistence within organisms and their capacity to induce lasting physiological changes, suggest that their effects may extend beyond the initial exposure period. Consequently, prior exposure to NP, or other emerging contaminants, may substantially alter the interactive effects of subsequent environmental stressors, yet this possibility has received little attention.

Here, we test if prior exposure to environmental NP generates carryover effects that modify subsequent responses to multiple environmental stressors. Using the blue mussel *Mytilus edulis* (Linnaeus 1758), a widespread foundation species in temperate coastal systems (Norling & Kautsky, 2007; Sendra et al., 2021), we experimentally examined how NP pre-exposure influences responses to subsequent copper contamination, nutrient enrichment, and reduced salinity, individually and in combination. Specifically, we asked whether previous NP exposure (1) alters organismal performance, metabolic rates and energy status following exposure to subsequent stressors, as these traits underpin stress tolerance and mediate trade-offs among maintenance, detoxification, and performance (Sokolova, 2013), and (2) can modify the nature of interactive effects among multiple stressors (*e.g.*, changing additive effects to synergistic or antagonistic outcomes). By incorporating environmental history into a multiple stressor framework, this study advances our understanding of how previous environmental experiences shape responses to environmental change and highlights the importance of temporal dynamics for predicting ecological responses in increasingly variable environments (Lundsgaard et al., 2023).

## 2. Materials and Methods

### 2.1. Experimental Design

We employed a factorial experimental design to assess how nanoplastic (NP) pre-exposure (absence [NP−] or presence [NP+]) influences the tolerance of *M. edulis* to three subsequent environmental stressors: nutrient enrichment ([N−] or [N+]), copper contamination ([Cu−] or [Cu+]), and freshwater input ([FW−] or [FW+]). Mussel assemblages, pre-exposed to NP or not, were subjected to all possible stressor combinations. This resulted in 16 treatment groups, with four replicates *per* treatment, totaling 64 experimental units (individual plates with mussel assemblages).

### 2.2. Mussel Collection

Adult blue mussels, *M. edulis*, were hand-collected over three days from floating pontoons at Malahide Marina in the Malahide Estuary on the east coast of Ireland (53°27′12″N, 6°9′5″W; Fig. S1a). This estuarine site experiences tidal ranges of 2–4 m and salinity fluctuations between 18 and 34 (Brooks et al., 2023). A total of 550 mussels (5–8 cm shell length) were randomly distributed among ten 45-L plastic containers used as water baths (five for NP pre-exposure and five controls), with 55 mussels per container. Each bath contained 30 L of seawater and was suspended from the pontoons so that approximately three-quarters of the volume of the bath remained submerged, thus maintaining ambient water temperature. Containers were continuously aerated, and mussels were acclimated for one week before NP pre-exposure under the same conditions.

### 2.3. NP Pre-Exposure

NP pre-exposure lasted six days and was conducted from the 15^th^ to 21^st^ of August 2023 (Fig. S1b). To expose mussels to NP, we followed a standardized protocol adapted from Blancho et al. (2021) to simulate environmentally relevant conditions. Briefly, 400 g of plastic debris (<10 cm diameter), including polyethylene (PE), polypropylene (PP), and polystyrene (PS), were combined with 600 mL of deionized water in a 1-L square bottle and stirred at 250 rpm for 48 h, followed by 1 h of sonication. Microplastics were removed using a 40-μm mesh filter, and residual organic material was degraded using 1% (wt%) hydrogen peroxide (H₂O₂) under ultraviolet (UV) light for 5 h (H₂O₂/UV). The remaining NP fraction was collected with a 1.2-μm glass fiber filter (Prat DUMAS). Final NP concentration in the stock solution was 0.15 g L⁻¹ (see below for NP characterization).

To administer the NP treatment, the NP stock was mixed daily with Shellfish Diet 1800 (Reed Mariculture, Campbell, CA, USA), used as the daily food source. For each 45-L container, 33 mL of Shellfish Diet (0.6 mL per mussel) was diluted in 330 mL of seawater. NP-exposed treatments received 20 mL of NP solution, while controls received 20 mL of deionized water to maintain consistent dilution volumes. Mussels filtered the feed in 190.9 ± 2.73 min (mean ± SE) on average, with filtration times of 192.4 ± 3.79 min under NP exposure and 189.5 ± 3.97 min in controls. Filtration was considered complete when the water became clear and mussel valves visibly closed. Temperature, oxygen, and salinity were measured daily in each water bath (Table S1) using a multiprobe (Temperature and Dissolved oxygen, Hach Lange LDO101, Düsseldorf, Germany) and salinometer (WTW Cond 3110, Weilheim, Germany). After the 6-day NP pre-exposure period, four mussels from both control and NP-exposed groups were collected, dissected for whole tissues, and freeze-dried in 10 mL glass vials for subsequent NP characterization.

### 2.4. Experimental System and Procedure

Following NP pre-exposure, experimental exposure to nutrient enrichment, copper contamination, and freshwater input lasted 21 days (21 August–11 September 2023; Fig. S1b). To assess the effects of multiple stressors on blue mussel assemblage tolerance, metabolism, and energy status, standardized mussel assemblages were created by securing eight mussels to concrete backing plates (29 × 25 cm) using soft gardening mesh (15 × 15 cm) and cable ties (Fig. S2). Plates were suspended from wooden buffer timbers at Malahide Marina, with their center positioned 25 cm below the water surface.

Nutrient enrichment was applied following the protocol of Worm et al. (2000). Polyethylene mesh diffuser bags (800 × 50 mm; 1-mm² mesh size) were filled with slow-release fertilizer pellets (Worm et al., 2000). Each bag was divided into four sections with cable ties, each containing 80 g of pellets, producing a nutrient release rate of 1 g cm⁻² to ensure consistent diffusion throughout the experiment. Bags were attached to the backing plates using cable ties to promote even nutrient distribution in the surrounding water column (Brooks & Crowe, 2018). Copper contamination and freshwater input were applied using a programmable dosing system following Browne et al. (2016). Six 640-L containers (three for copper, three for freshwater) were fitted with pumps and digital timers to regulate delivery. Copper concentration was maintained at 425 µg L⁻¹, consistent with previous studies (Brooks & Crowe, 2019; Browne et al., 2016). Copper was mixed with seawater and continuously delivered to the experimental plates. For the freshwater treatment, freshwater was delivered to the plates four times daily for 2 h each (08:00–10:00; 14:00–16:00; 20:00–22:00; 02:00–04:00). Plates were randomly assigned to one of 16 treatment groups and spaced 1.5 m apart along the dock to minimize interactions among experimental units (Brooks et al., 2023; Brooks & Crowe, 2019; Browne et al., 2016; see also Table 1). Environmental parameters, including temperature, oxygen, and salinity, were measured daily at 09:30 near ten experimental plates during freshwater input (five exposed and five unexposed; Table S1). Different plates were sampled each day.

### 2.5. Nanoplastic Characterization in Mussels

Freeze-dried mussel tissues were analyzed for nanoplastics using pyrolysis coupled with gas chromatography–mass spectrometry (Py-GC/MS), following Blancho et al. (2021). Briefly, tissues were chemically digested, extracted, and analyzed to identify and quantify polystyrene (PS), polypropylene (PP), and polyethylene (PE). Detailed extraction procedures, analytical settings, calibration methods, and data processing are provided in Appendix S1, and results are shown in Figure S3.

### 2.6. Characterization of Community Metabolism

Community oxygen absorption rates were measured as a proxy for their metabolic rates (Fry & Hart, 1948) by monitoring changes in dissolved oxygen concentrations over time. Experimental plates containing live mussel assemblages were placed in 6-L dark incubation chambers equipped with a pump to ensure water circulation and homogeneous conditions. Dissolved oxygen concentrations were measured at the start of the incubation and after 10, 20, and 30 min using an optical oxygen probe (HQ20, Hach Lange Ltd., Loveland, USA). Respiration rates were calculated from the linear decline in oxygen concentration between 10 and 30 min, allowing an initial acclimation period and ensuring that photosynthesis had ceased (Noël et al., 2010). The measurement interval was selected based on previous linearity tests conducted under the same experimental conditions (Nielsen, 2001). Respiration rates were standardized *per* hour, normalized to the number of live mussels *per* plate, and expressed as mg O₂ L⁻¹ h⁻¹ *per* live mussel (Brooks et al., 2023; Brooks & Crowe, 2019).

### 2.7. Survival

After measuring community oxygen absorption rates, all organisms were removed from the experimental plates. Mortality was assessed as the percentage of dead mussels relative to the initial number *per* plate, based on the presence of dead individuals and empty shells.

### 2.8. Determination of Mussels Energy Status

ATP content was measured individually in the abductor muscle of four mussels *per* experimental plate as a proxy for energy status. Half of each abductor muscle was flash-frozen in liquid nitrogen, stored at –80°C, and analyzed using a luminescence-based ATP detection assay following Chida and Kido (2014) with minor modifications. ATP concentrations were quantified from calibration curves and standardized to the total wet mass of each abductor muscle. Detailed extraction procedures, assay conditions, and quality control are provided in Appendix S2.

### 2.9. Statistical Analyses

All data were grouped into three response variables: community oxygen absorption rates, survival, and ATP content. The effects of NP pre-exposure, freshwater input (Salinity), nutrient enrichment (Nutrient), and copper contamination (Copper) were analyzed using generalized linear models (GLMs) or generalized linear mixed models (GLMMs), depending on the response variable. Community oxygen absorption rates were analyzed using a Gamma GLM with a log-link function, survival using a binomial GLM with a logit-link function, and ATP content using a Gamma GLMM with a log-link function. Plate was included as a random effect for ATP content based on likelihood ratio tests, and mussel length was retained as a covariate because it significantly improved model fit. Candidate models of increasing complexity were compared using likelihood ratio tests (LRTs) and Akaike’s Information Criterion (AIC) to identify the most parsimonious model for each response. The final models retained the main effects, selected two-way interactions, and a Nanoplastics × Nutrient × Copper three-way interaction for community oxygen absorption rates, main effects only for survival, and the main effects plus a Salinity × Nutrient interaction for ATP content. When interaction terms were retained, post hoc pairwise comparisons were performed using estimated marginal means implemented in the emmeans package (Lenth, 2025).

Model assumptions were assessed using residual diagnostics, overdispersion tests, and Cook’s distance. Predictor significance was evaluated using Type III Wald χ² tests (Gamma models) or Type III likelihood-ratio χ² tests (binomial GLM). To quantify treatment effects relative to controls, we calculated Hedges’ *g* effect sizes, allowing standardized comparisons while correcting for small sample sizes (negligible: |*g*| < 0.2; small: 0.2 ≤ |*g*| < 0.5; medium: 0.5 ≤ |*g*| < 0.8; large: |*g*| ≥ 0.8). Analyses were conducted in R (version 4.4.1) using the glmmTMB (Brooks et al., 2017), lme4 (Bates et al., 2015), **car** (Fox & Weisberg, 2019), and DHARMa R packages. Full model outputs are provided in Table S2.

### 2.10. Classification of interactive effects

When significant interactions were detected, we applied the interaction classification framework of Crain et al. (2008), later expanded by Piggott et al. (2015), using Hedges’ g effect sizes to distinguish additive, antagonistic, and synergistic responses (Crain et al., 2008; Piggott et al., 2015). We differentiated between the combined stressor effect size, representing the overall biological response to simultaneous stressor exposure relative to controls, and the interaction effect size, representing deviations from additive expectations. Additive effects were inferred when the 95% confidence intervals of the Hedges’ g interaction effect overlapped zero, indicating no deviation from additivity (Gurevitch et al., 2000). Synergistic effects occurred when the observed combined effect exceeded additive expectations, whereas antagonistic effects occurred when the combined effect was lower than expected.

## 3. Results

### 3.1. Community Metabolism

Community oxygen absorption rates averaged 0.065 ± 0.004 mg (mean ± SE) O₂ L⁻¹ h⁻¹ mussel⁻¹ across treatments, ranging from 0.029 ± 0.009 mg under control conditions to 0.119 ± 0.034 mg in mussel communities pre-exposed to nanoplastics (NP) and subsequently exposed to combined copper contamination and nutrient enrichment (Table S3, Fig. S4). Community respiration was influenced by a significant three-way interaction among NP pre-exposure, copper contamination, and nutrient enrichment (χ^2^ = 6.99, df = 1, *P* = 0.0082; Table S2, Fig. 1a). In the absence of NP pre-exposure, oxygen absorption rates under combined nutrient enrichment and copper contamination did not differ from control conditions, whereas NP pre-exposure resulted in significantly higher oxygen absorption rates under the same combined stressors (Fig. 1a, Table S4). This response exceeded additive expectations, indicating a positive synergistic interactive effect (Fig. 1b). Relative to controls, nearly all stressor combinations produced large effect sizes, except for the Cu+:Nu+ treatment without NP pre-exposure, which showed only a small effect (Table S5, Fig. S5).

**Figure 1.**
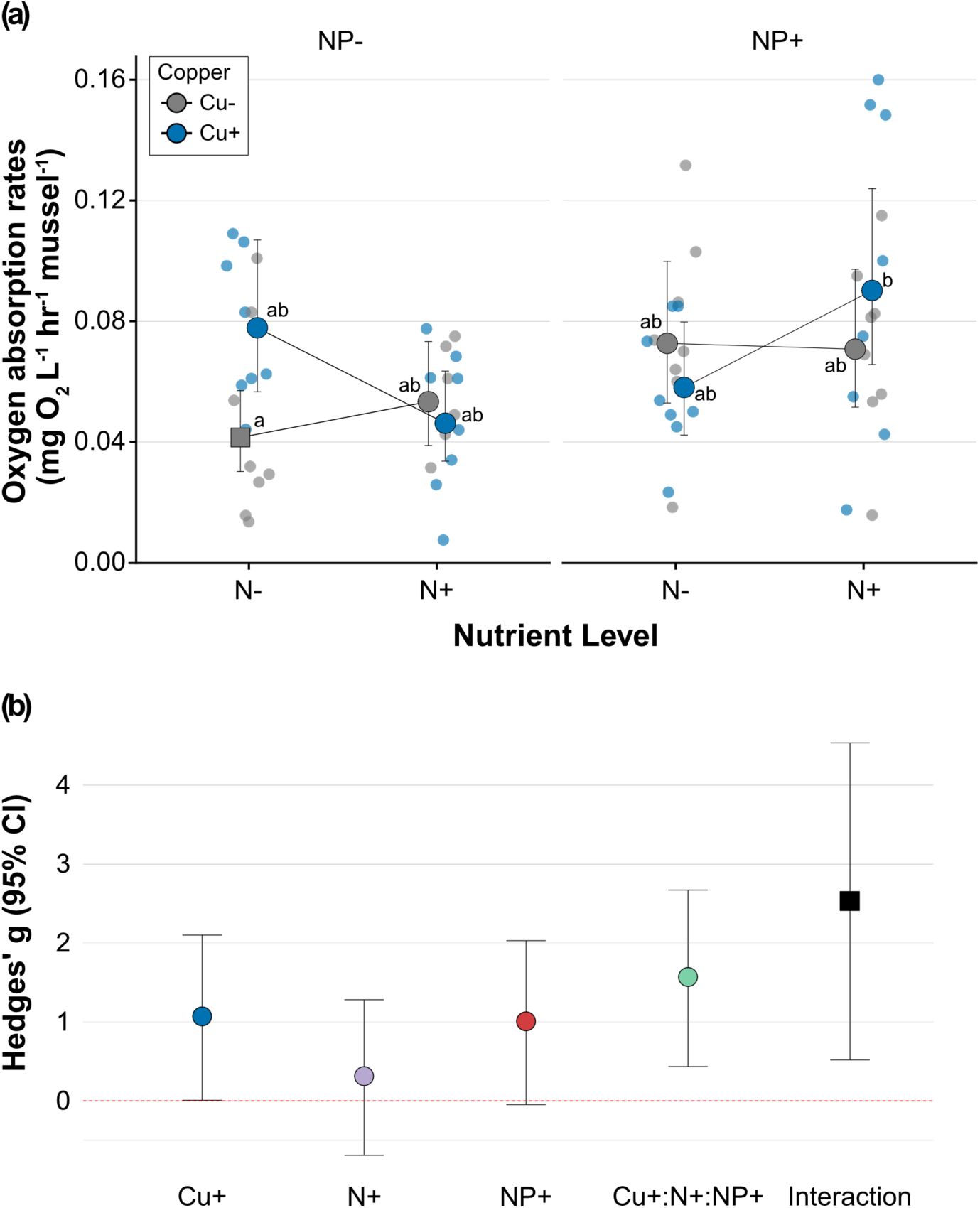
Interactive effects of nanoplastic (NP) pre-exposure, copper (Cu), and nutrient enrichment (N) on community respiration in *M. edulis*. **(a)** Interaction between copper and nutrients on respiration in mussels with and without NP pre-exposure. Different lowercase letters indicate significant differences among treatment combinations. The full results of the pairwise comparisons are presented in Supplementary Table S4. The control treatment is represented by a square, and error bars indicate 95% confidence intervals. **(b)** Hedge’s *g* effect sizes (±95% CI) showing the individual and combined impacts of NP pre-exposure, copper, and nutrient enrichment on community respiration relative to the control conditions. The interaction effect size (i.e. the deviation from the additive expectation) is represented by a black square. Interactive effects are considered significant when the interaction effect size and its 95% confidence interval do not overlap zero.

### 3.2. Survival

Survival averaged 61.0 ± 2.3% (mean ± SE) across treatments, ranging from 96.9 ± 3.1% under control conditions to 40.6 ± 3.1% in mussels pre-exposed to NP and subsequently exposed to combined copper contamination and nutrient enrichment (Table S6, Fig. S6). NP pre-exposure significantly reduced survival regardless of subsequent stressor exposure (χ^2^_1_ = 17.49, df = 1, *P* < 0.001; Table S3, Fig. 2). Freshwater input and copper contamination also reduced survival by 14.9% (χ^2^ = 4.66, df = 1, *P* = 0.031) and 25.3% (χ^2^ = 19.03, df = 1, *P* < 0.001), respectively (Table S2, Fig. 2). Hedges’g indicated large, directionally negative effect sizes for all individual and combined stressor treatments relative to controls (Table S7, Fig. S7).

**Figure 2.**
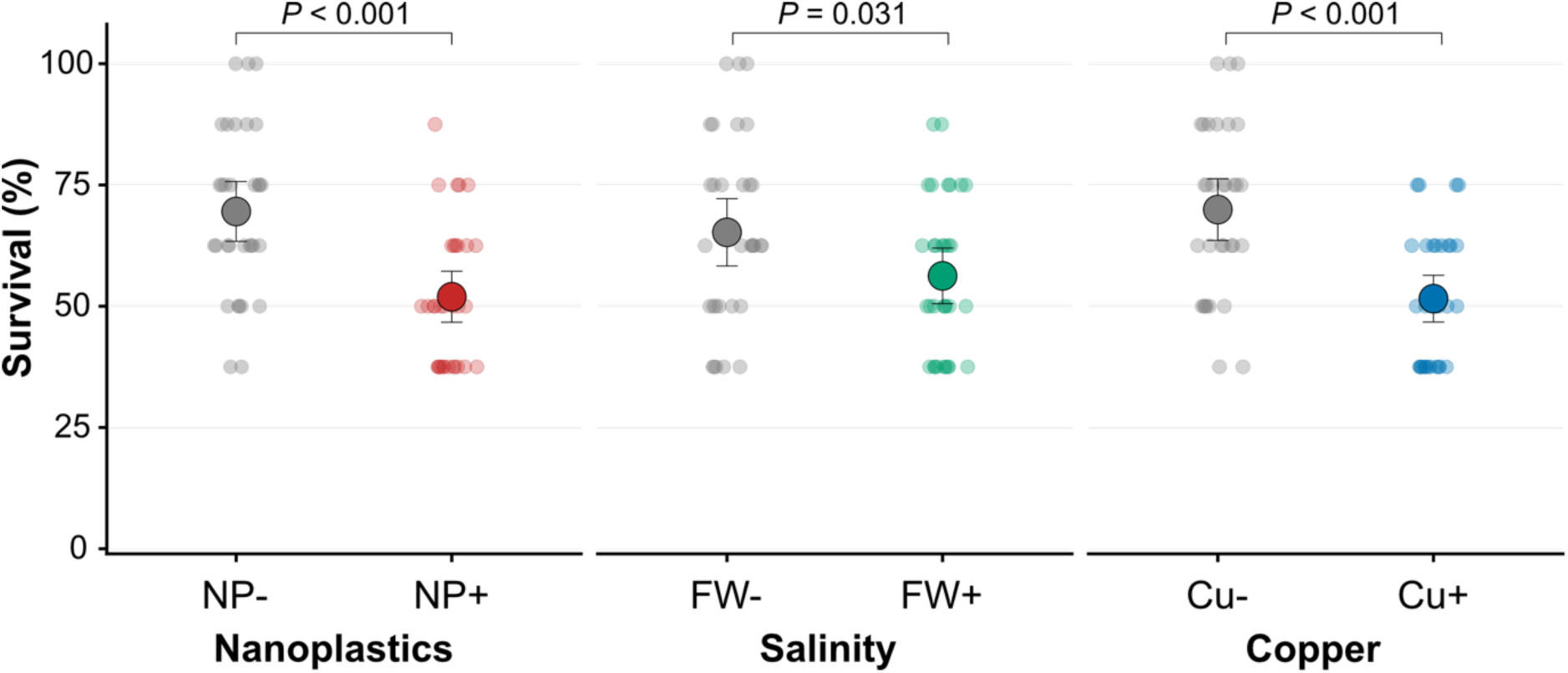
Main effects of nanoplastic pre-exposure, salinity and copper on survival in *Mytilus edulis*. Raw observations are shown as individual points, and large circles represent model-estimated marginal means (±95% CI). The displayed *P*-value corresponds to the Type III likelihood ratio test for the main effect of salinity. Full model results are provided in Supplementary Table S2.

### 3.3. ATP Content

ATP content in *M. edulis* abductor muscles averaged 2.94 ± 0.27 µg (mean ± SE) across treatments, ranging from 5.59 ± 1.99 µg in mussels pre-exposed to NP to 1.25 ± 0.16 µg in NP pre-exposed mussels subjected to freshwater input (Table S8, Fig. S8). ATP content was influenced by a significant interaction between freshwater input and nutrient enrichment (χ^2^ = 6.45, df = 1, *P* = 0.011, Table S2, Fig. 3a). In the absence of nutrient enrichment, freshwater input significantly reduced ATP content, whereas this effect was no longer evident under nutrient-enriched conditions (Fig. 3a, Table S9). Consequently, the combined effect of freshwater input and nutrient enrichment was less negative than expected under additivity, indicating an antagonistic interactive effect (Fig. 3b). Across all treatments, only the N+:FW+ and NP+:N+:FW+ combinations produced slight increases in ATP relative to controls, although these effects were negligible. All other treatments reduced ATP content compared to control conditions, with effect sizes ranging from negligible to large (Table S10, Fig. S9).

**Figure 3.**
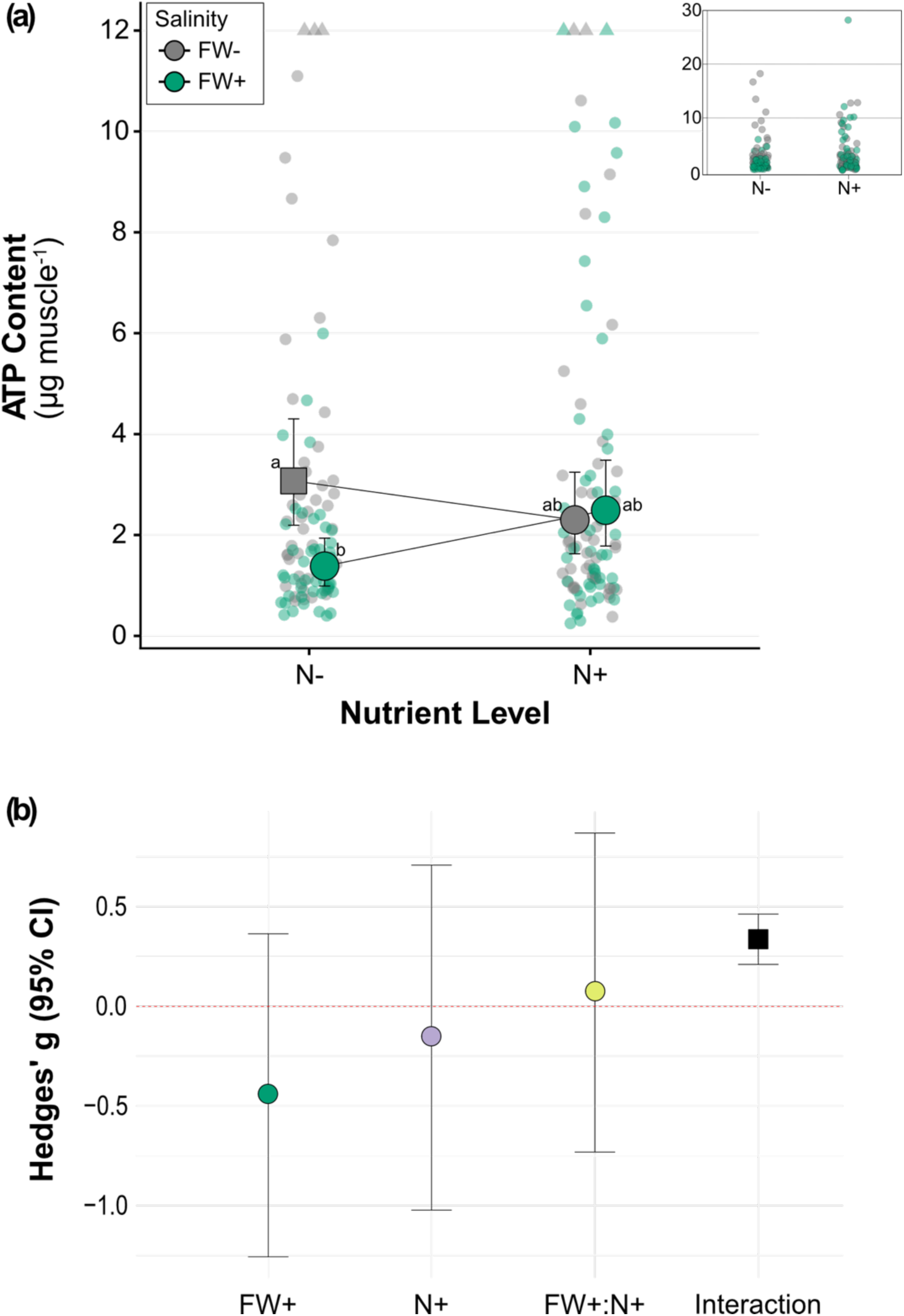
Interactive effects of nutrient enrichment (N) and salinity (FW) on ATP content in *M. edulis*. **(a)** Interaction between nutrient enrichment and freshwater input on ATP content. Different lowercase letters indicate significant differences among treatment combinations. The full results of the pairwise comparisons are presented in Supplementary Table S9. The control treatment is represented by a square, and error bars indicate 95% confidence intervals. **(b)** Hedge’s *g* effect sizes (±95% CI) showing the individual and combined impacts of freshwater input and nutrient enrichment on ATP content relative to the control conditions. The interaction effect size (i.e. the deviation from the additive expectation) is represented by a black square. Interactive effects are considered significant when the interaction effect size and its 95% confidence interval do not overlap zero.

## 4. Discussion

Ecological legacies arising from previous environmental conditions are increasingly recognized as important determinants of ecological responses to global changes (Antonelli et al. 2024), yet they remain largely overlooked in multiple stressor research. Our results suggest that environmental history should be considered as a fundamental dimension in multiple stressor ecology. Specifically, prior nanoplastic (NP) exposure reduces survival regardless of subsequent stressor exposure and modify the interactive effects of copper contamination and nutrient enrichment on community metabolic costs, demonstrating that even short-term environmental experiences can leave persistent biological legacies. We first discuss the role of carryover effects as a temporal dimension of multiple stressor ecology, before considering the underlying mechanisms that may generate these responses, the context dependence of cross-tolerance, and finally the implications of incorporating dynamic temporal stress regimes into future multiple stressor research.

### 4.1 Environmental history shapes responses to multiple stressors

An important implication of our findings is that the ecological consequences of multiple stressors cannot always be inferred from present-day environmental conditions alone. Rather, organisms exposed to identical stressor combinations can respond differently depending on their environmental history (Harrison et al., 2011; Brooks & Crowe, 2019; Antonelli et al., 2024). In our study, previous nanoplastic (NP) exposure consistently reduce survival regardless of subsequent stressor exposure and created interactive effects between copper contamination and nutrient enrichment on community metabolic rates, with a positive synergistic response emerging only in previously exposed mussels.

Although carryover effects have been documented across a broad range of environmental contexts (Harrison et al., 2011; Brooks & Crowe, 2019; Rodgers et al., 2025), they have rarely been explicitly incorporated into a multiple stressor framework. For example, sequential exposure to warming and hypoxia in oysters generated context-dependent carryover effects, whereby the consequences of previous environmental conditions depended on the subsequent stress environment (Donelan et al., 2021). Despite growing evidence that previous environmental conditions influence later organismal performance, most multiple stressor studies continue to focus on simultaneous rather than sequential exposures (Jackson et al., 2021). Within the context of plastic pollution, research has similarly concentrated on interactions between NP and concurrent environmental stressors (*e.g.,* ocean acidification, Manno et al., 2022; temperature, salinity, and natural particulate matter, Lins et al., 2022), whereas comparatively little attention has been given to whether prior NP exposure leaves biological legacies that influence responses after exposure has ceased. Our findings extend previous work by demonstrating that contaminant exposure can generate biological legacies that persist beyond the initial exposure period.

Importantly, environmental history operates across multiple temporal scales. Carryover effects may arise from relatively brief exposures that induce persistent physiological changes within an individual, as observed here, but they may also emerge following repeated or chronic exposure or be transmitted across generations through parental effects and transgenerational plasticity (Harrison et al., 2011; Donelan et al., 2021; Rodgers et al., 2025). Although these processes differ in their underlying mechanisms and persistence, they share a common feature: previous environmental conditions influence future ecological responses. Viewing environmental history as a temporal continuum provides an opportunity to integrate traditionally separate fields—including carryover effects, ecological legacies, parental effects, and transgenerational plasticity—within a unified framework for multiple stressor ecology.

An important outcome of our study is that the effects of NP pre-exposure differed among biological responses. Previous NP exposure modified the interactive effects of multiple stressors on community metabolic rates, whereas survival reflects a persistent carryover effect that is largely independent of subsequent environmental conditions. This pattern suggests that different biological responses integrate previous environmental experiences over different temporal scales, with some reflecting persistent physiological impairment and others becoming evident only following subsequent environmental challenges.

### 4.2 Mechanisms linking environmental history to ecological responses

Environmental legacies ultimately manifest through organismal, population, community, and ecosystem responses, but they originate from physiological and cellular processes within individuals. Across diverse environmental contexts, prior conditions have been shown to alter subsequent survival, growth, and reproduction through persistent physiological changes (Hettinger et al. 2012; Donelan et al. 2021). The durability of these changes provides the mechanistic basis through which environmental history shapes future ecological outcomes, with consequences that may propagate upward to population dynamics, species interactions, and ecosystem functioning.

Nanoplastics provide one additional example of how physiological legacies may arise from environmental history. Previous work shows that NP and microplastics alter clearance rates, immune function, oxidative balance, and energy allocation in bivalves (Paul-Pont et al., 2016; Magni et al., 2018; Sendra et al., 2020, 2021). Importantly, physiological impairments may persist beyond the exposure period, even when depuration appears complete (Elizalde-Velázquez et al. 2020; Sun et al. 2022, 2023). This persistence provides a plausible mechanism through which prior NP exposure influences subsequent responses in our study, demonstrating how contaminants can leave lasting physiological signatures that shape later ecological performance.

The magnitude and persistence of these ecological legacies depend strongly on the characteristics of the initial exposure. Stressor sequence, intensity, duration, frequency, and recovery time influence the extent to which physiological disruption is repaired or persists through time (Hettinger et al. 2012, 2013). Likewise, in the context of NP pollution, particle size, polymer type, and surface chemistry are also likely to influence the magnitude and persistence of these physiological disruptions (Ward et al., 2019; Sendra et al., 2021). Identifying which characteristics of contaminant exposure drive persistent physiological changes – and over what temporal scales – will be essential for predicting how environmental history influences tolerance to future stress. Together, these exposure-dependent physiological legacies form the foundation upon which higher-level ecological responses emerge, but translating individual disruption into population or community-level outcomes remains a major challenge.

Extending from individual physiology to higher-order ecological processes, predicting how these legacies scale remains difficult. Although some approaches assume that ecosystem responses can be predicted by scaling individual responses, this scaling is unlikely to be linear because physiological changes may be buffered, amplified, or modified by demographic processes, species interactions, and food-web dynamics (Smith et al. 2019; Enquist et al. 2003). Our results illustrate this complexity, with prior NP exposure modifying community-level responses while producing a persistent reduction in survival independent of subsequent environmental conditions.

### 4.3 Context-dependent responses to multiple stressors

Research on multiple stressors has traditionally emphasized their negative, often synergistic, effects on organismal performance and ecosystem functioning (Côté et al., 2016). However, antagonistic interactions are also common and may arise when one stressor partially offsets the physiological consequences of another (Rodgers et al. 2025). In our study, the antagonistic interaction between nutrient enrichment and freshwater input illustrates how physiological responses can modify the outcome of multiple stressor interactions independently of environmental history. Whereas freshwater input reduced mussels’ energy status under ambient nutrient conditions, this effect disappeared under nutrient enrichment, suggesting that nutrient availability partially compensated for the energetic demands imposed by osmotic stress. *Mytilus* spp. are osmoconformers capable to tolerate reductions in salinity through intracellular volume regulation rather than active osmoregulation (Davenport & Redpath, 1984; Gosling, 2003; Hauton, 2016). One possible explanation is that increased nutrient availability enhanced the energetic capacity (Josefson & Rasmussen, 2000), ultimately supporting a greater ability to maintain cellular homeostasis during osmotic stress. Similar antagonistic interactions between nutrient enrichment and freshwater input have previously been reported in *Mytilus* spp. and *Macoma balthica*, where nutrient enrichment partially buffered the biological consequences of reduced salinity (Carrier-Belleau et al., 2021). Together, these findings suggest that improved energetic status may partially offset the physiological costs of other stress (Thomsen et al. 2013). The outcome of multiple stressor exposure therefore depends not only on stressor identity, but also on the physiological state through which organisms experience and respond to environmental change.

### 4.4 Toward a temporally explicit multiple stressor ecology

Our findings suggest that environmental history should be explicitly incorporated into experimental designs and ecological predictions. We propose three complementary directions for advancing a temporally explicit framework for multiple stressor ecology.

First, future studies should explicitly incorporate previous environmental history into experimental designs. Environmental history can be represented experimentally through sequential exposure designs, preconditioning experiments, trans- and multigenerational or reciprocal exposure approaches that distinguish current from previous environmental conditions. Alternatively, naturally contrasting populations with different histories of contaminant exposure, thermal variability, hypoxia, or other environmental pressures provide an opportunity to evaluate how long-term environmental experience influences subsequent stress tolerance. Extending this perspective across generations through parental exposure and transgenerational plasticity experiments would further integrate ecological and evolutionary responses within a common temporal framework (Vermandele et al. 2025; Chakravarti et al. 2016; Donelson et al. 2017).

Second, greater attention should be given to the temporal characteristics of stress exposure itself. Importantly, these temporal characteristics should be considered relative to the biological timescales of the focal organisms (*i.e.*, life cycle and generation time), as the same stress event may represent a short-term transient perturbation for one species but persist across much or all of the life cycle of another (Saether et al. 2005). Experimental approaches capable of explicitly quantifying these temporal dimensions—including exposure gradients and response-surface designs spanning multiple stressor intensities, exposure durations, and recovery periods (*e.g.*, Thomas & Ranjan, 2024)—offer a promising avenue for identifying thresholds beyond which physiological recovery does no longer occur and ecological legacies emerge.

Third, understanding ecological legacies requires integrating responses across multiple levels of biological organization. Carryover effects originate through cellular and physiological processes but may only become fully apparent at higher levels of biological organization through changes in individual performance, population dynamics, species interactions, community structure, and ecosystem functioning. Consequently, reliance on a single biological response is unlikely to capture the full ecological consequences of previous environmental experience. Integrative, multi-level frameworks combining physiological, behavioral, demographic, community, and ecosystem responses (Calosi et al., 2013; Madeira et al., 2024) offer a promising direction for linking mechanistic processes to ecological outcomes and improving predictions of biological responses under multiple stressors.

Taken together, our findings demonstrate that previous environmental conditions fundamentally shape responses to multiple stressors. Explicitly incorporating environmental history into experiments and predictive frameworks will improve our ability to predict ecological responses in increasingly dynamic environments.

## 5. Authors contributions

**Charlotte Carrier-Belleau:** conceptualization, data curation, formal analysis, investigation, methodology, project administration, funding acquisition, validation, visualization, writing - original draft, writing - review and editing. **Paul Brooks:** conceptualization, methodology, funding acquisition, resources, writing - review and editing. **Lauric Feugere:** methodology, formal analysis (lead for ATP analysis), writing - original draft, writing - review and editing. **Philippe Archambault:** conceptualization, writing - review and editing. **Piero Calosi:** conceptualization, methodology, funding acquisition, writing - review and editing. **Jennifer Coughlan:** conceptualization, methodology, writing - review and editing. **Alexina Trudelle:** methodology, formal analysis, writing - original draft, writing - review and editing. **Tasman Crowe:** conceptualization, methodology, funding acquisition, resources, writing - review and editing, **Julien Gigault:** conceptualization, methodology, formal analysis, funding acquisition, writing - review and editing.

## 6. Funding

C.C.B. was supported by a postdoctoral fellowship from the Fonds de Recherche du Québec – Nature et technologies (grant ID 314292), the Sentinel North Postdoctoral Fellowship, and the Joint Arctic Postdoctoral Fellowship Awards Program from MEOPAR and ArcticNet. PC was supported by an NSERC Discovery grant (RGPIN-2020-05627) and a Canada Foundation for Innovation (CFI) grant. LF acknowledges support from Mitacs Accelerate towards PC [IT35557] and funding by Fonds de recherche du Québec – Nature et technologies [PBEEE #350373].

## 7. Acknowledgements

We thank M. Kirwan, A. Murphy, C. Udusalu, R. Rahill and T. Eardley for their assistance during the experiment, and J.J. Piggott for his help and advice on multistressor interactions. We thank the staff at Malahide Marina for access, storage, and facilities to do this experiment. The authors wish to thank Jonathan Coudé for help during ATP determination.

## Notes

### Competing Interest Statement

The authors have declared no competing interest.

