## Supplementary Material for "From Bad to Worse: Carryover Effects Alter Responses to Multiple Stressors"

*Ecology*

**Appendix S1.** Detailed methods for nanoplastic extraction and characterization

To prepare samples for NP characterization, freeze-dried mussel tissues were crushed using a mortar. For each sample, 2 mL of 0.1% potassium hydroxide (KOH) was added and agitated for 24 h. Samples were then centrifuged at 4000 rpm for 10 min (25°C; Centrifuge 5702, Eppendorf Co.), after which 1 mL of supernatant was transferred to a separate glass vial. A dichloromethane extraction was subsequently performed on the supernatant, an efficient method for extracting NP from environmental samples, particularly water samples with high organic matter content. Recovery rates reached up to 70%, which is considered acceptable based on similar published methods (Okoffo & Thomas, 2024).

Extracted particles were identified and quantified using Pyrolysis coupled to Gas Chromatography and Mass Spectrometry (Py-GC/MS; 5977B Inert Plus MSD with Intuvo 9000 GC, Agilent Technologies, Inc.), following established methods (Blancho et al., 2021). Data processing was conducted using Qualitative Analysis Navigator (Version 10.0, Agilent Technologies, Inc.) and MS Quantitative Analysis (Quant-My-Way) (Version B.09.00, Agilent Technologies, Inc.). Pyrolyzates were identified by comparing mass spectra with the NIST library using NIST MS Search program 2.3 (NIST, Gaithersburg, MD, USA). We focused on polystyrene, polypropylene, and polyethylene, as these are among the most widely produced and used plastics. Nanoplastic quantification was based on internal calibration curves for polystyrene and polypropylene NP using standard mixtures, with quantifiers at $m/z = 104$ for polystyrene and $m/z = 70$ for polypropylene. Py-GC/MS detected alkanes and alkenes that may originate from both natural organic matter and nanometric polyethylene (chain fragments up to C20). Given the higher detection limit for PE compared to PP (~200 ng vs. 10–20 ng) and the absence of alkadienes, these signals could not be unequivocally assigned to PE. Relative alkene abundances normalized to deuterated polystyrene showed slight differences between exposed and unexposed organisms (Fig. S2). Overall, the absence of polypropylene markers and alkadienes prevents definitive conclusions regarding PE presence in organisms at the end of exposure, suggesting partial or total NP excretion by mussels.

**Appendix S2.** Detailed methods for ATP extraction and characterization

To assess mussel energy status, we measured individually the ATP content of their abductor muscle. Four mussels *per* plate were carefully dissected, and half of each abductor muscle was flash-frozen in liquid nitrogen before being stored at –80°C at University College Dublin (Ireland). Samples were then shipped frozen in a dry shipper to the University of Quebec in Rimouski (Canada), where ATP analyses were performed.

ATP was extracted following the methods developed by Chida and Kido (2014). We validated the optimal wet tissue mass range as ~30 mg and the optimal ATP extract dilution as 10-fold, by comparing different dilutions of a pool of all 187 experimental samples (undiluted, and 10 and 100-time diluted samples in deionized water). Prior to ATP extraction, all abductor muscle samples were randomized to be processed blind to the experimenter and prevent batch effects. ATP was extracted in batches of 20 samples, for a total of 187 abductor muscle samples. Entire abductor muscle samples were thawed, quickly blotted on absorbent paper to remove excess water, and weighed (242.33 ± 5.68 mg) on an analytical scale (#VWR-225AC, VWR International, Radnor, PA, USA) to obtain a given mussel's total abductor muscle wet mass. Next, ~30 mg (31.38 ± 0.21 mg) subsamples were cut with fine scissors and weighed on a plastic vessel using the same analytical scale. When not handled, microcentrifuge tubes were always kept on ice to prevent overheating. The 30-mg subsamples were transferred to a reinforced microcentrifuge tube (#15-340-162, Thermo Fisher Scientific, Waltham, MA, USA) containing 1 mL of pre-chilled phenol-TE (#P4557, Sigma-Aldrich, St. Louis, MO, USA) and six ceramic beads (2.8 mm, #15-340-160, Thermo Fisher Scientific, Waltham, MA, USA). Subsamples were homogenized using the Fast-prep homogenizer (Fast-prep®-24.

MP Biomedicals, Irvine, CA, USA) at 5 m $s^{-1}$ for two cycles of 30 s interspaced by a 30 s-cooling of microcentrifuge tubes in ice to prevent overheating. A volume of 0.8 mL of the homogenized supernatant was then transferred to a new microcentrifuge tube containing 200 µL of pre-chilled chloroform (#C2432, Sigma-Aldrich, St. Louis, MO, USA) and 150 µL of pre-chilled deionized water. Homogenates were vigorously shaken for 20 s, quickly shaken and inverted to ensure optimal mixing. Homogenates were centrifuged for 5 min at 10,000 g at room temperature in a Mini-Spin® centrifuge (Eppendorf, Hamburg, Germany). A volume of 150 µL of the upper aqueous phase was carefully transferred using a micropipette to a new microcentrifuge tube, ensuring the interphase was not disturbed. These resulting concentrated ATP extracts were stored at -80 ºC until further processing. ATP contents were quantified using a standard luminescence-based ATP detection kit (ATP Detection Assay Kit - luminescence, #700515, Cayman chemical, Ann Arbor, MI, USA) following the manufacturer's recommendations. Briefly, 10 µL of ATP extract samples were diluted 10-fold in 90 µL of deionized water. Luminescence was read using a plate reader (Agilent Biotek Energy H1 Synergy H1, Agilent Technologies. Santa Clara, CA, USA) accompanied by the BioTek Gen5 3.12 software (Agilent Technologies. Santa Clara, CA, USA; 10-s orbital shaking at 237 cpm, 4 mm, one-read only, integration time: 1 s, delay: 100 ms, gain: 240). ATP was quantified in triplicate across 8 batches of 96-well plate runs and with the aid of a multichannel micropipette. Coefficients of variation between triplicates were 5.17 ± 0.26 %, validating the suitability of our data (e.g., Drew & Leeuwenburgh, 2003). For each plate, duplicate eight-point standard curves (all with $R^2 > 0.98$) were used to determine, for each plate well, ATP contents in nM from measured log-transformed blank-corrected relative luminescence units, following the manufacturer's recommendations. ATP contents from 10 μL of a given sample in the plate well were reverted back to the amount of ATP in the entire abductor muscle wet mass of a given individual mussel.

**Table S1**. Mean values (± SE) of experimental parameters measured in water baths during the pre-exposure phase or near experimental plates during the exposure phase. Temperature, salinity, and dissolved oxygen were measured daily in all 10 water baths under control (NP−) and NP-exposed (NP+) conditions, and near 10 experimental plates in the absence (FW−) and presence (FW+) of freshwater input.

| **Pre-exposure phase** | | | |
|---|---|---|---|
| Treatment | Temperature (ºC) | Salinity (PSU) | Dissolved oxygen (%) |
| NP- | 18.92 ± 0.11 | 31.29 ± 0.12 | 90.10 ± 0.47 |
| NP+ | 19.29 ± 0.13 | 31.26 ± 0.11 | 89.12 ± 0.48 |
| **Exposure phase** | | | |
| Treatment | Temperature (ºC) | Salinity (PSU) | Dissolved oxygen (%) |
| FW- | 21.30 ± 0.09 | 31.20 ± 0.07 | 96.90 ± 0.12 |
| FW+ | 21.39 ± 0.09 | 23.35 ± 0.10 | 96.89 ± 0.14 |

**Table S2.** Complete Type III analysis of variance (ANOVA) results for the effects of nanoplastic pre-exposure, freshwater input ("Salinity"), nutrient enrichment ("Nutrient"), and copper contamination ("Copper") on community oxygen absorption rates, survival, and ATP content in *Mytilus edulis*. Community respiration was analysed using a model including a three-way interaction, survival using a model with main effects only, and ATP content using a model including two-way interactions.

**Community respiration rates**

| Factor | Df | $\chi^2$ | *p*-value |
|---|---|---|---|
| (Intercept) | 1 | 299.3578 | <0.001 |
| Salinity | 1 | 3.0318 | 0.0816 |
| Nanoplastic | 1 | 5.0392 | 0.0249 |
| Nutrient | 1 | 0.8023 | 0.3704 |
| Copper | 1 | 6.0274 | 0.0141 |
| Nanoplastics:Nutrient | 1 | 0.5814 | 0.4458 |
| Nanoplastics:Copper | 1 | 5.8994 | 0.0151 |
| Nutrient:Copper | 1 | 4.5470 | 0.0330 |
| Nanoplastics:Nutrient:Copper | 1 | 6.9905 | 0.0082 |

**Survival**

| Factor | Df | $\chi^2$ | *p*-value |
|---|---|---|---|
| Nanoplastics | 1 | 17.4901 | <0.001 |
| Salinity | 1 | 4.6612 | 0.0309 |
| Nutrient | 1 | 0.2218 | 0.6377 |
| Copper | 1 | 19.0325 | <0.001 |

**ATP content**

| Factor | Df | $\chi^2$ | *p*-value |
|---|---|---|---|
| (Intercept) | 1 | 0.0144 | 0.9045 |
| Nanoplastics | 1 | 0.0567 | 0.8117 |
| Salinity | 1 | 10.7526 | 0.0010 |
| Nutrient | 1 | 1.3938 | 0.2378 |
| Copper | 1 | 2.9857 | 0.0840 |
| Length | 1 | 5.7274 | 0.0167 |
| Salinity:Nutrient | 1 | 6.4585 | 0.0110 |

**Table S3.** Mean community oxygen absorption rates (mg of $O_2$ $L^{-1}$ $h^{-1}$ mussel $alive^{-1}$) in the blue mussel *Mytilus edulis* across experimental treatments: nanoplastics (NP), copper contamination (Cu), nutrient enrichment (N) and salinity variation (FW). Results are shown for the number of replicates (n), the mean, standard error (SE) and 95% Confidence Interval (CI_Lower and CI_Upper).

| Treatment | Nanoplastic | Copper | Nutrient | Salinity | n | Mean | SE | CI_Lower | CI_Upper |
|---|---|---|---|---|---|---|---|---|---|
| 1 | NP- | Cu- | N- | FW- | 4 | 0.0287 | 0.0092 | -0.0009 | 0.0583 |
| 2 | NP- | Cu+ | N- | FW- | 4 | 0.0747 | 0.0139 | 0.0304 | 0.1190 |
| 3 | NP- | Cu- | N+ | FW- | 4 | 0.0471 | 0.0065 | 0.0265 | 0.0677 |
| 4 | NP- | Cu- | N- | FW+ | 4 | 0.0599 | 0.0188 | 0.0000 | 0.1198 |
| 5 | NP- | Cu+ | N+ | FW- | 4 | 0.0367 | 0.0112 | 0.0009 | 0.0725 |
| 6 | NP- | Cu+ | N- | FW+ | 4 | 0.0811 | 0.0124 | 0.0418 | 0.1204 |
| 7 | NP- | Cu- | N+ | FW+ | 4 | 0.0603 | 0.0076 | 0.0363 | 0.0844 |
| 8 | NP- | Cu+ | N+ | FW+ | 4 | 0.0582 | 0.0113 | 0.0222 | 0.0941 |
| 9 | NP+ | Cu- | N- | FW- | 4 | 0.0540 | 0.0122 | 0.0151 | 0.0930 |
| 10 | NP+ | Cu+ | N- | FW- | 4 | 0.0584 | 0.0090 | 0.0297 | 0.0872 |
| 11 | NP+ | Cu- | N+ | FW- | 4 | 0.0656 | 0.0174 | 0.0100 | 0.1211 |
| 12 | NP+ | Cu- | N- | FW+ | 4 | 0.0977 | 0.0132 | 0.0558 | 0.1396 |
| 13 | NP+ | Cu+ | N+ | FW- | 4 | 0.1194 | 0.0340 | 0.0110 | 0.2277 |
| 14 | NP+ | Cu+ | N- | FW+ | 4 | 0.0577 | 0.0137 | 0.0141 | 0.1012 |
| 15 | NP+ | Cu- | N+ | FW+ | 4 | 0.0764 | 0.0143 | 0.0307 | 0.1220 |
| 16 | NP+ | Cu+ | N+ | FW+ | 4 | 0.0681 | 0.0126 | 0.0282 | 0.1081 |

**Table S4.** Tukey-adjusted pairwise comparisons of estimated marginal means for the Nanoplastic x Nutrient x Copper interaction on community respiration rates. Ratios are presented on the response scale, and *P*-values were adjusted using Tukey's method.

| Treatment 1 | Treatment 2 | Ratio | SE | df | *z* ratio | *p*-value |
|---|---|---|---|---|---|---|
| NP−/N−/Cu− | NP+/N−/Cu− | 0.5800 | 0.1407 | Inf | -2.2448 | 0.3248 |
| NP−/N−/Cu− | NP−/N+/Cu− | 0.8044 | 0.1955 | Inf | -0.8957 | 0.9865 |
| NP−/N−/Cu− | NP+/N+/Cu− | 0.6061 | 0.1474 | Inf | -2.0587 | 0.4421 |
| NP−/N−/Cu− | NP−/N−/Cu+ | 0.5502 | 0.1339 | Inf | -2.4551 | 0.2151 |
| NP−/N−/Cu− | NP+/N−/Cu+ | 0.7342 | 0.1789 | Inf | -1.2679 | 0.9107 |
| NP−/N−/Cu− | NP−/N+/Cu+ | 0.9217 | 0.2237 | Inf | -0.3359 | 1.0000 |
| NP−/N−/Cu− | NP+/N+/Cu+ | 0.4420 | 0.1086 | Inf | -3.3234 | 0.0201 |
| NP+/N−/Cu− | NP−/N+/Cu− | 1.3868 | 0.3367 | Inf | 1.3465 | 0.8809 |
| NP+/N−/Cu− | NP+/N+/Cu− | 1.0449 | 0.2539 | Inf | 0.1808 | 1.0000 |
| NP+/N−/Cu− | NP−/N−/Cu+ | 0.9485 | 0.2306 | Inf | -0.2177 | 1.0000 |
| NP+/N−/Cu− | NP+/N−/Cu+ | 1.2658 | 0.3080 | Inf | 0.9689 | 0.9788 |
| NP+/N−/Cu− | NP−/N+/Cu+ | 1.5890 | 0.3855 | Inf | 1.9088 | 0.5447 |
| NP+/N−/Cu− | NP+/N+/Cu+ | 0.7621 | 0.1867 | Inf | -1.1089 | 0.9551 |
| NP−/N+/Cu− | NP+/N+/Cu− | 0.7535 | 0.1828 | Inf | -1.1667 | 0.9413 |
| NP−/N+/Cu− | NP−/N−/Cu+ | 0.6839 | 0.1660 | Inf | -1.5656 | 0.7710 |
| NP−/N+/Cu− | NP+/N−/Cu+ | 0.9128 | 0.2216 | Inf | -0.3759 | 1.0000 |
| NP−/N+/Cu− | NP−/N+/Cu+ | 1.1459 | 0.2781 | Inf | 0.5611 | 0.9993 |
| NP−/N+/Cu− | NP+/N+/Cu+ | 0.5496 | 0.1340 | Inf | -2.4559 | 0.2147 |
| NP+/N+/Cu− | NP−/N−/Cu+ | 0.9077 | 0.2202 | Inf | -0.3991 | 0.9999 |
| NP+/N+/Cu− | NP+/N−/Cu+ | 1.2114 | 0.2940 | Inf | 0.7905 | 0.9936 |
| NP+/N+/Cu− | NP−/N+/Cu+ | 1.5208 | 0.3692 | Inf | 1.7267 | 0.6697 |
| NP+/N+/Cu− | NP+/N+/Cu+ | 0.7293 | 0.1776 | Inf | -1.2962 | 0.9006 |
| NP−/N−/Cu+ | NP+/N−/Cu+ | 1.3346 | 0.3238 | Inf | 1.1897 | 0.9351 |
| NP−/N−/Cu+ | NP−/N+/Cu+ | 1.6754 | 0.4069 | Inf | 2.1247 | 0.3987 |
| NP−/N−/Cu+ | NP+/N+/Cu+ | 0.8035 | 0.1955 | Inf | -0.8992 | 0.9862 |
| NP+/N−/Cu+ | NP−/N+/Cu+ | 1.2553 | 0.3051 | Inf | 0.9357 | 0.9826 |
| NP+/N−/Cu+ | NP+/N+/Cu+ | 0.6021 | 0.1464 | Inf | -2.0872 | 0.4232 |
| NP−/N+/Cu+ | NP+/N+/Cu+ | 0.4796 | 0.1173 | Inf | -3.0056 | 0.0538 |

**Table S5**. Standardized effect sizes (Hedges'g) comparing community respiration rates under nanoplastics (NP), copper contamination (Cu), nutrient enrichment (N) and salinity variation (FW) treatments to unstressed control condition (Treatment 1), with 95% confidence intervals and Hedges'g magnitude thresholds (Large: $|g| \geq 0.8$).

| Treatment | Nanoplastic | Copper | Nutrient | Salinity | Hedges'g | CI_Lower | CI_Upper | Magnitude |
|---|---|---|---|---|---|---|---|---|
| 2 | NP- | Cu+ | N- | FW- | 1.69 | 0.07 | 3.30 | Large |
| 3 | NP- | Cu- | N+ | FW- | 1.00 | -0.47 | 2.47 | Large |
| 4 | NP- | Cu- | N- | FW+ | 0.92 | -0.54 | 2.37 | Large |
| 5 | NP- | Cu+ | N+ | FW- | 0.34 | -1.06 | 1.73 | Small |
| 6 | NP- | Cu+ | N- | FW+ | 2.08 | 0.36 | 3.80 | Large |
| 7 | NP- | Cu- | N+ | FW+ | 1.62 | 0.03 | 3.22 | Large |
| 8 | NP- | Cu+ | N+ | FW+ | 1.24 | -0.27 | 2.75 | Large |
| 9 | NP+ | Cu- | N- | FW- | 1.01 | -0.46 | 2.48 | Large |
| 10 | NP+ | Cu+ | N- | FW- | 1.41 | -0.14 | 2.96 | Large |
| 11 | NP+ | Cu- | N+ | FW- | 1.15 | -0.35 | 2.64 | Large |
| 12 | NP+ | Cu- | N- | FW+ | 2.63 | 0.74 | 4.53 | Large |
| 13 | NP+ | Cu+ | N+ | FW- | 1.58 | 0.01 | 3.17 | Large |
| 14 | NP+ | Cu+ | N- | FW+ | 1.08 | -0.41 | 2.56 | Large |
| 15 | NP+ | Cu- | N+ | FW+ | 1.71 | 0.09 | 3.33 | Large |
| 16 | NP+ | Cu+ | N+ | FW+ | 1.55 | -0.03 | 3.13 | Large |

**Table S6**. Mean survival rates (%) of *M. edulis* across experimental treatments: nanoplastics (NP), copper contamination (Cu), nutrient enrichment (N) and salinity variation (FW). Results are shown for the number of replicates (n), the mean, standard error (SE) and 95% Confidence Interval (CI_Lower and CI_Upper).

| Treatment | Nanoplastic | Copper | Nutrient | Salinity | n | Mean | SE | CI_Lower | CI_Upper |
|---|---|---|---|---|---|---|---|---|---|
| 1 | NP- | Cu- | N- | FW- | 4 | 96.88 | 3.125 | 86.930 | 106.820 |
| 2 | NP- | Cu+ | N- | FW- | 4 | 71.88 | 3.125 | 61.930 | 81.820 |
| 3 | NP- | Cu- | N+ | FW- | 4 | 78.12 | 5.984 | 59.082 | 97.169 |
| 4 | NP- | Cu- | N- | FW+ | 4 | 75.00 | 5.103 | 58.760 | 91.240 |
| 5 | NP- | Cu+ | N+ | FW- | 4 | 62.50 | 5.103 | 46.260 | 78.740 |
| 6 | NP- | Cu+ | N- | FW+ | 4 | 50.00 | 5.103 | 33.760 | 66.240 |
| 7 | NP- | Cu- | N+ | FW+ | 4 | 68.75 | 8.069 | 43.072 | 94.428 |
| 8 | NP- | Cu+ | N+ | FW+ | 4 | 56.25 | 8.069 | 30.572 | 81.928 |
| 9 | NP+ | Cu- | N- | FW- | 4 | 65.62 | 5.984 | 46.582 | 84.669 |
| 10 | NP+ | Cu+ | N- | FW- | 4 | 43.75 | 3.608 | 32.266 | 55.234 |
| 11 | NP+ | Cu- | N+ | FW- | 4 | 65.62 | 7.864 | 40.597 | 90.653 |
| 12 | NP+ | Cu- | N- | FW+ | 4 | 50.00 | 5.103 | 33.760 | 66.240 |
| 13 | NP+ | Cu+ | N+ | FW- | 4 | 40.62 | 3.125 | 30.680 | 50.570 |
| 14 | NP+ | Cu+ | N- | FW+ | 4 | 43.75 | 6.250 | 23.860 | 63.640 |
| 15 | NP+ | Cu- | N+ | FW+ | 4 | 56.38 | 9.375 | 29.540 | 89.210 |
| 16 | NP+ | Cu+ | N+ | FW+ | 4 | 46.88 | 5.984 | 27.832 | 65.919 |

**Table S7**: Standardized effect sizes (Hedges'g) comparing survival under nanoplastics (NP), copper contamination (Cu), nutrient enrichment (N) and salinity variation (FW) treatments to unstressed control condition (Treatment 1), with 95% confidence intervals and Hedges'g magnitude thresholds (Large: $|g| \geq 0.8$).

| Treatment | Nanoplastic | Copper | Nutrient | Salinity | Hedges'g | CI_Lower | CI_Upper | Magnitude |
|---|---|---|---|---|---|---|---|---|
| 2 | NP- | Cu+ | N- | FW- | -3.62 | -5.87 | -1.37 | Large |
| 3 | NP- | Cu- | N+ | FW- | -1.71 | -3.33 | -0.09 | Large |
| 4 | NP- | Cu- | N- | FW+ | -2.25 | -4.02 | -0.48 | Large |
| 5 | NP- | Cu+ | N+ | FW- | -3.53 | -5.75 | -1.31 | Large |
| 6 | NP- | Cu+ | N- | FW+ | -4.82 | -7.55 | -2.08 | Large |
| 7 | NP- | Cu- | N+ | FW+ | -2.00 | -3.70 | -0.30 | Large |
| 8 | NP- | Cu+ | N+ | FW+ | -2.89 | -4.87 | -0.91 | Large |
| 9 | NP+ | Cu- | N- | FW- | -2.85 | -4.81 | -0.88 | Large |
| 10 | NP+ | Cu+ | N- | FW- | -6.84 | -10.47 | -3.21 | Large |
| 11 | NP+ | Cu- | N+ | FW- | -2.27 | -4.05 | -0.49 | Large |
| 12 | NP+ | Cu- | N- | FW+ | -4.82 | -7.55 | -2.08 | Large |
| 13 | NP+ | Cu+ | N+ | FW- | -7.83 | -11.90 | -3.75 | Large |
| 14 | NP+ | Cu+ | N- | FW+ | -4.67 | -7.35 | -2.00 | Large |
| 15 | NP+ | Cu- | N+ | FW+ | -2.33 | -4.13 | -0.54 | Large |
| 16 | NP+ | Cu+ | N+ | FW+ | -4.55 | -7.78 | -1.93 | Large |

**Table S8**: Mean ATP concentration (µg per abductor mussel) in *Mytilus edulis* across experimental treatments: nanoplastics (NP), copper contamination (Cu), nutrient enrichment (N) and salinity variation (FW). Results are shown for the number of replicates (n), the mean, standard error (SE) and 95% Confidence Interval (CI_Lower and CI_Upper).

| Treatment | Nanoplastic | Copper | Nutrient | Salinity | n | Mean | SE | CI_Lower | CI_Upper |
|---|---|---|---|---|---|---|---|---|---|
| 1 | NP- | Cu- | N- | FW- | 12 | 4.81 | 1.43 | 1.66 | 7.95 |
| 2 | NP- | Cu+ | N- | FW- | 12 | 2.74 | 0.49 | 1.66 | 3.82 |
| 3 | NP- | Cu- | N+ | FW- | 9 | 3.95 | 1.37 | 0.79 | 7.11 |
| 4 | NP- | Cu- | N- | FW+ | 12 | 2.38 | 0.44 | 1.42 | 3.34 |
| 5 | NP- | Cu+ | N+ | FW- | 13 | 2.41 | 0.41 | 1.52 | 3.30 |
| 6 | NP- | Cu+ | N- | FW+ | 11 | 1.28 | 0.37 | 0.47 | 2.10 |
| 7 | NP- | Cu- | N+ | FW+ | 12 | 5.18 | 2.36 | -0.01 | 10.37 |
| 8 | NP- | Cu+ | N+ | FW+ | 12 | 1.89 | 0.33 | 1.17 | 2.61 |
| 9 | NP+ | Cu- | N- | FW- | 10 | 5.59 | 1.99 | 1.10 | 10.09 |
| 10 | NP+ | Cu+ | N- | FW- | 12 | 2.29 | 0.35 | 1.52 | 3.06 |
| 11 | NP+ | Cu- | N+ | FW- | 12 | 4.19 | 1.23 | 1.49 | 6.90 |
| 12 | NP+ | Cu- | N- | FW+ | 12 | 1.25 | 0.16 | 0.89 | 1.61 |
| 13 | NP+ | Cu+ | N+ | FW- | 12 | 1.34 | 0.17 | 0.97 | 1.71 |
| 14 | NP+ | Cu+ | N- | FW+ | 12 | 1.27 | 0.19 | 0.85 | 1.70 |
| 15 | NP+ | Cu- | N+ | FW+ | 12 | 3.81 | 1.03 | 1.55 | 6.07 |
| 16 | NP+ | Cu+ | N+ | FW+ | 12 | 3.28 | 0.91 | 1.28 | 5.28 |

**Table S9**: Tukey-adjusted pairwise comparisons of estimated marginal means for the Salinity × Nutrient interaction on ATP concentration. Ratios are presented on the response scale, and *P*-values were adjusted using Tukey's method.

| Treatment 1 | Treatment 2 | Ratio | SE | df | *z* ratio | *p*-value |
|---|---|---|---|---|---|---|
| FW-/N- | FW+/N- | 2.2209 | 0.5404 | Inf | 3.2791 | 0.0057 |
| FW-/N- | FW-/N+ | 1.3365 | 0.3284 | Inf | 1.1806 | 0.6391 |
| FW-/N- | FW+/ N+ | 1.2332 | 0.2984 | Inf | 0.8664 | 0.8222 |
| FW+/ N- | FW-/N+ | 0.6018 | 0.1480 | Inf | -2.0643 | 0.1649 |
| FW+/ N- | FW+/N+ | 0.5553 | 0.1348 | Inf | -2.4239 | 0.0726 |
| FW-/N+ | FW+/ N+ | 0.9227 | 0.2261 | Inf | -0.3283 | 0.9878 |

**Table S10**: Standardized effect sizes (Hedges'g) comparing ATP production under nanoplastics (NP), copper contamination (Cu), nutrient enrichment (N) and salinity variation (FW) treatments to unstressed control condition (Treatment 1), with 95% confidence intervals and Hedges'g magnitude thresholds (Negligible: |g| <0.2, Small: 0.2 ≤ |g| < 0.5, Medium: 0.5 ≤ |g| < 0.8, Large: |g| ≥ 0.8).

| Treatment | Nanoplastic | Copper | Nutrient | Salinity | Hedges'g | CI_Lower | CI_Upper | Magnitude |
|---|---|---|---|---|---|---|---|---|
| 2 | NP- | Cu+ | N- | FW- | -0.54 | 1.35 | 0.28 | Medium |
| 3 | NP- | Cu- | N+ | FW- | -0.18 | 1.04 | 0.69 | Negligible |
| 4 | NP- | Cu- | N- | FW+ | -0.64 | 1.46 | 0.18 | Medium |
| 5 | NP- | Cu+ | N+ | FW- | -0.65 | 1.45 | 0.16 | Medium |
| 6 | NP- | Cu+ | N- | FW+ | -0.92 | 1.78 | 0.06 | Large |
| 7 | NP- | Cu- | N+ | FW+ | 0.05 | 0.75 | 0.85 | Negligible |
| 8 | NP- | Cu+ | N+ | FW+ | -0.79 | 1.62 | 0.05 | Medium |
| 9 | NP+ | Cu- | N- | FW- | 0.14 | 0.70 | 0.98 | Negligible |
| 10 | NP+ | Cu+ | N- | FW- | -0.67 | 1.50 | 0.15 | Medium |
| 11 | NP+ | Cu- | N+ | FW- | -0.13 | 0.93 | 0.67 | Negligible |
| 12 | NP+ | Cu- | N- | FW+ | -0.98 | 7.82 | 0.13 | Large |
| 13 | NP+ | Cu+ | N+ | FW- | -0.95 | 7.79 | 0.11 | Large |
| 14 | NP+ | Cu+ | N- | FW+ | -0.97 | 7.81 | 0.12 | Large |
| 15 | NP+ | Cu- | N+ | FW+ | -0.22 | 1.03 | 0.58 | Small |
| 16 | NP+ | Cu+ | N+ | FW+ | -0.36 | 1.16 | 0.45 | Small |

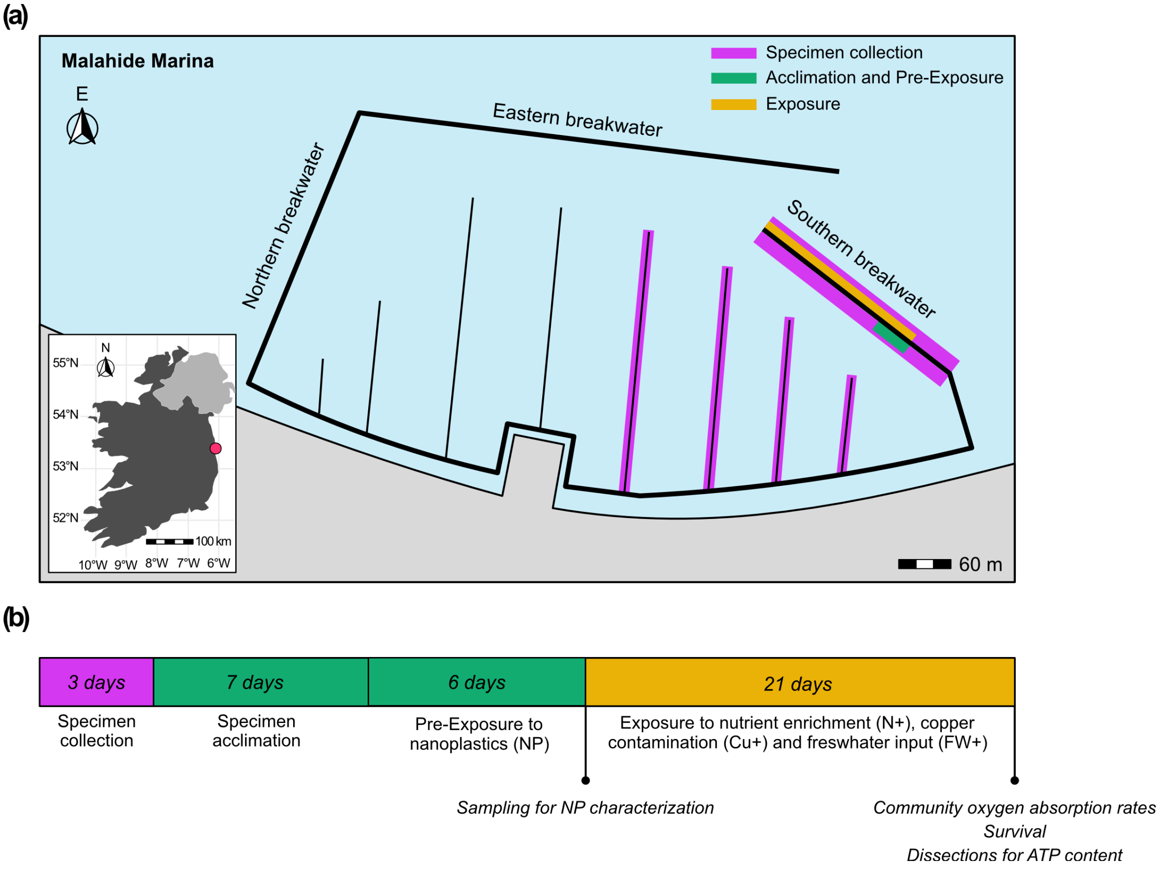


**Figure S1.** Schematic showing (a) location of experimental set-up and the distribution of experimental units during the pre-exposure phase to nanoplastics (NP; green) and the exposure phase (in orange) to copper ($Cu^{+}$), elevated nutrients ($N^{+}$), and freshwater input (FW+), and (b) the experimental timeline.

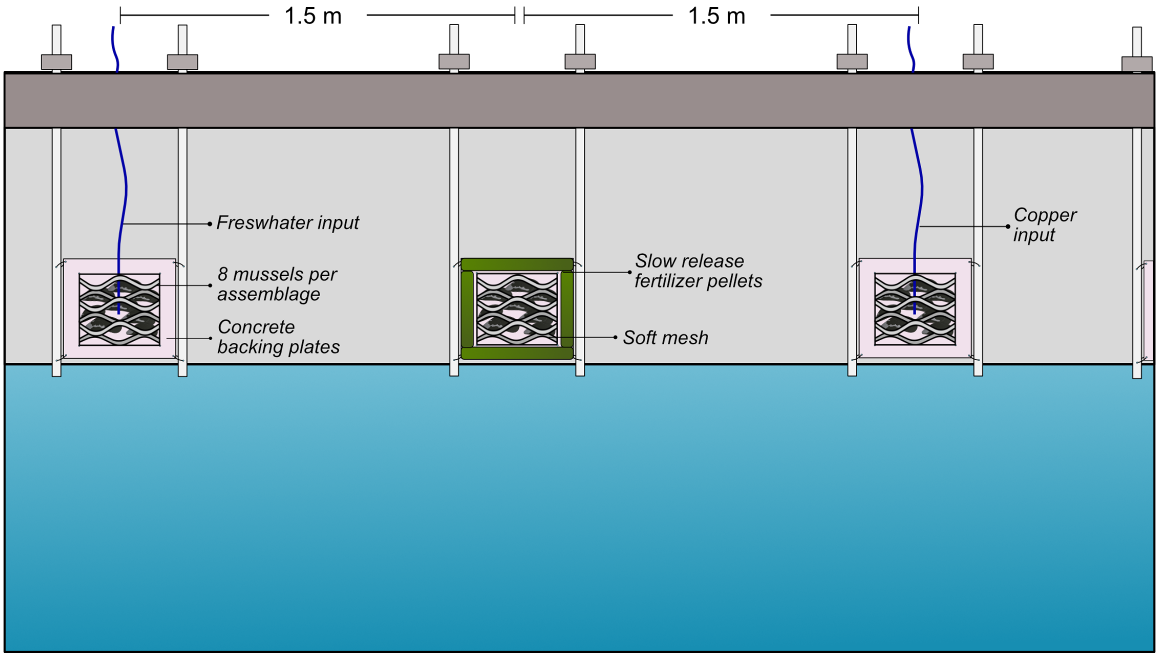


**Figure S2.** Schematic representation of the experimental setup of treatment plates. Each plate contained eight mussels secured to concrete backing plates with soft mesh. Nutrient enrichment was achieved using slow-release fertilizer pellets, while copper and freshwater inputs were delivered *via* tubing connected to header tanks.

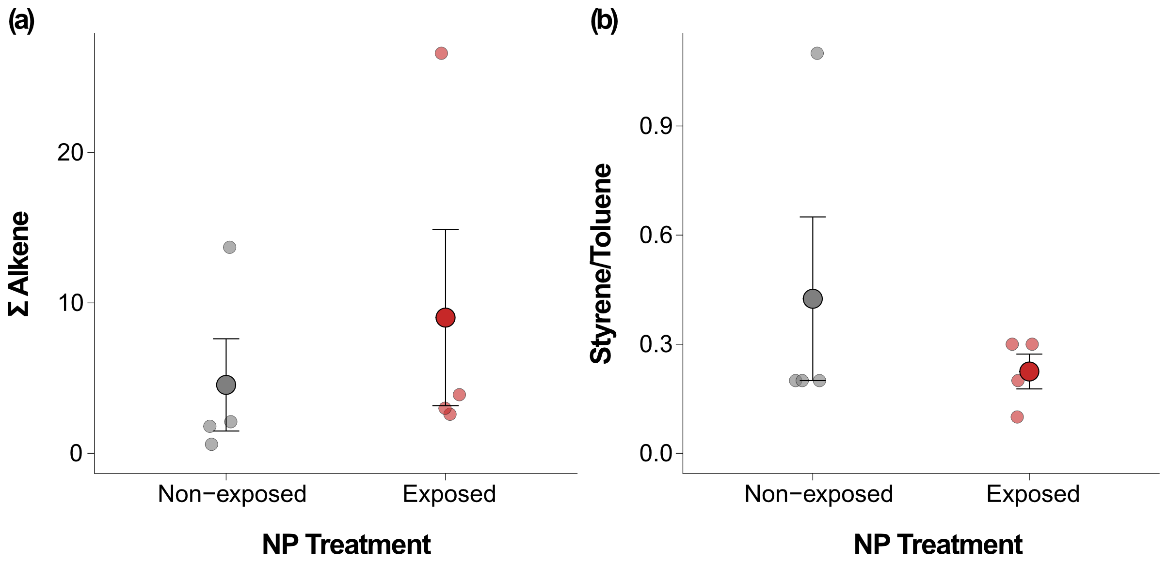


**Figure S3.** Nanoplastic characterization in mussel tissues from non-exposed and exposed organism (a) Total alkene signal extracted from the pyrograms at *m/z* 69. (b) Toluene-to-styrene ratio, calculated from pyrogram peak intensities at *m/z* 91 (toluene) and *m/z* 104 (styrene). Raw observations are shown as individual points, and large circles represent means ± SE.

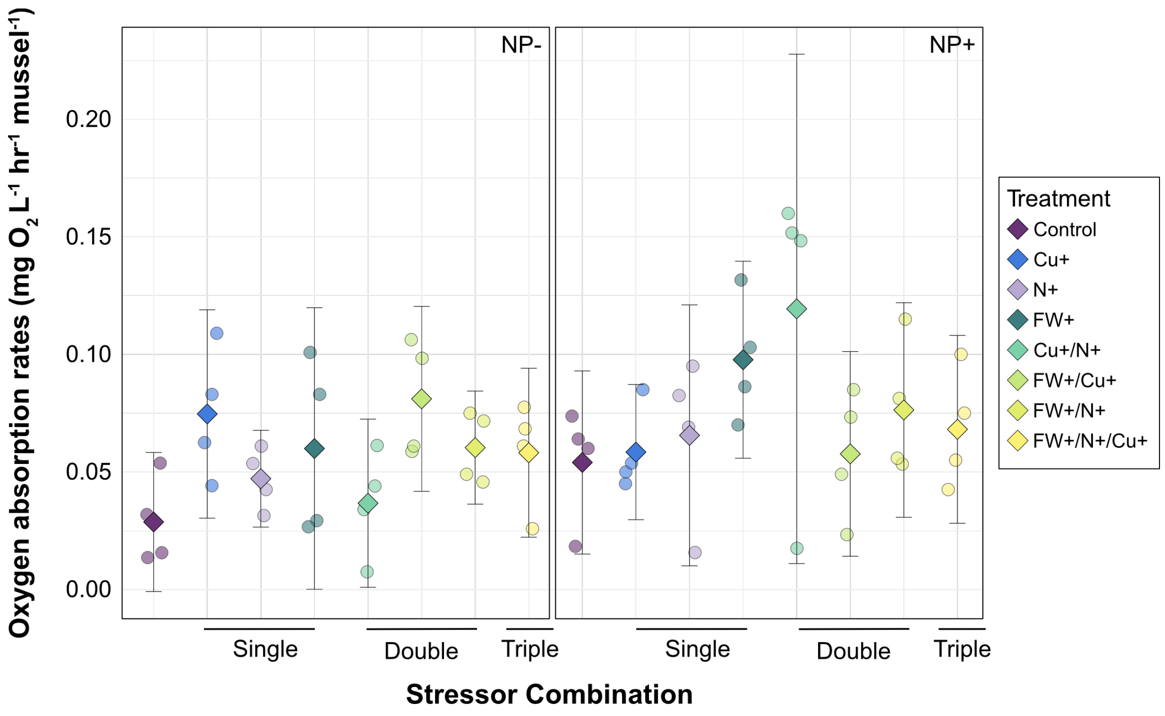


**Figure S4**. Mean community oxygen absorption rates across all experimental conditions. Results are presented for individual and combined stressors, including copper contamination ($Cu^{+}$), nutrient enrichment ($N^{+}$), and freshwater input (FW+), under conditions with and without nanoplastic pre-exposure ($NP^{+}$). Error bars represent 95% confidence intervals.

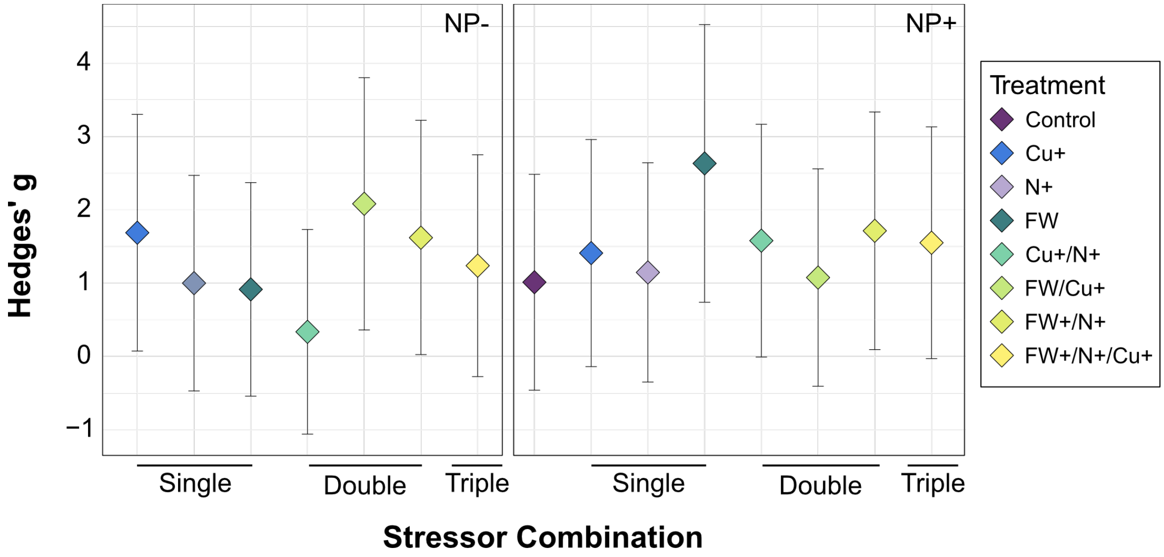


**Figure S5.** Standardized effect sizes (Hedges'g) comparing community respiration rates under individual and combined stressors, relative to control conditions (not shown on figure). Error bars correspond to 95% confidence intervals. Negative results show a decrease in survival rates compared to control conditions.

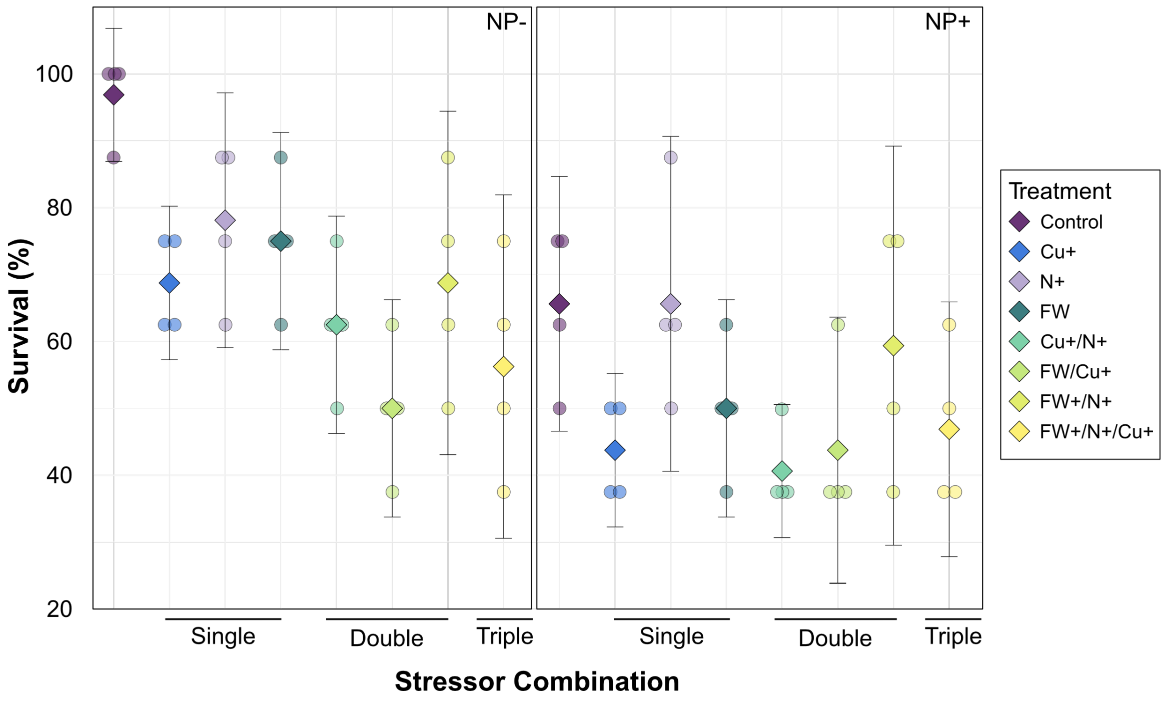


**Figure S6.** Mean survival rates (%) of *Mytilus edulis* across all experimental conditions. Results are presented for individual and combined stressors, including copper contamination ($Cu^+$), nutrient enrichment ($N^+$), and freshwater input (FW), under conditions with and without nanoplastic pre-exposure ($NP^+$). Error bars represent 95% confidence intervals.

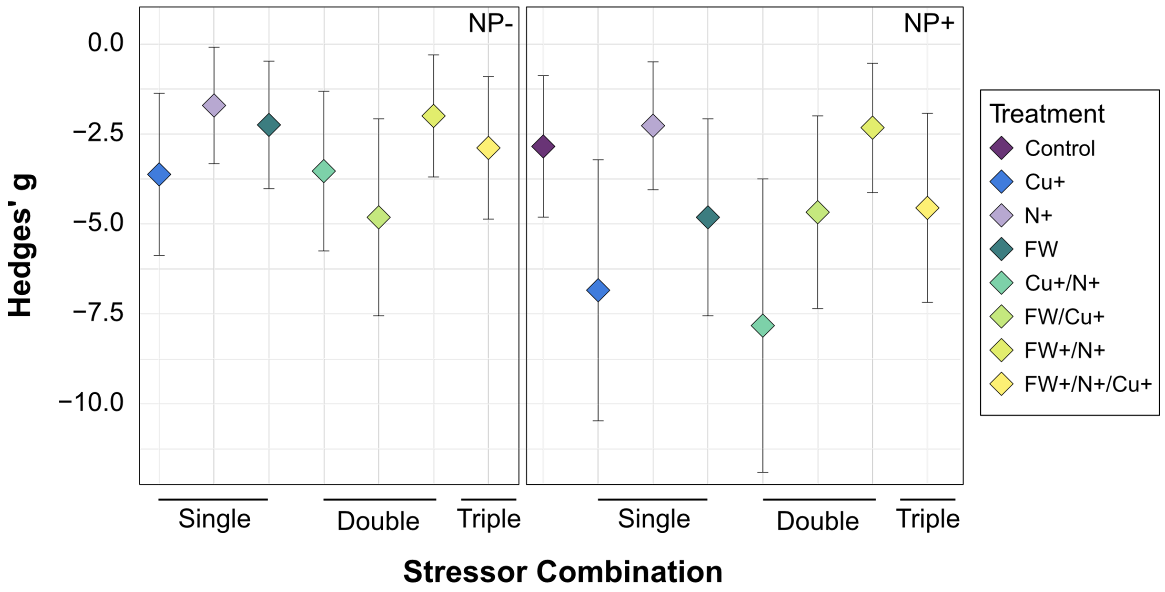


**Figure S7.** Standardized effect sizes (Hedges'g) comparing survival rates under individual and combined stressors, relative to control conditions (not shown on figure). Error bars correspond to 95% confidence intervals. Negative results show a decrease in survival rates compared to control conditions.

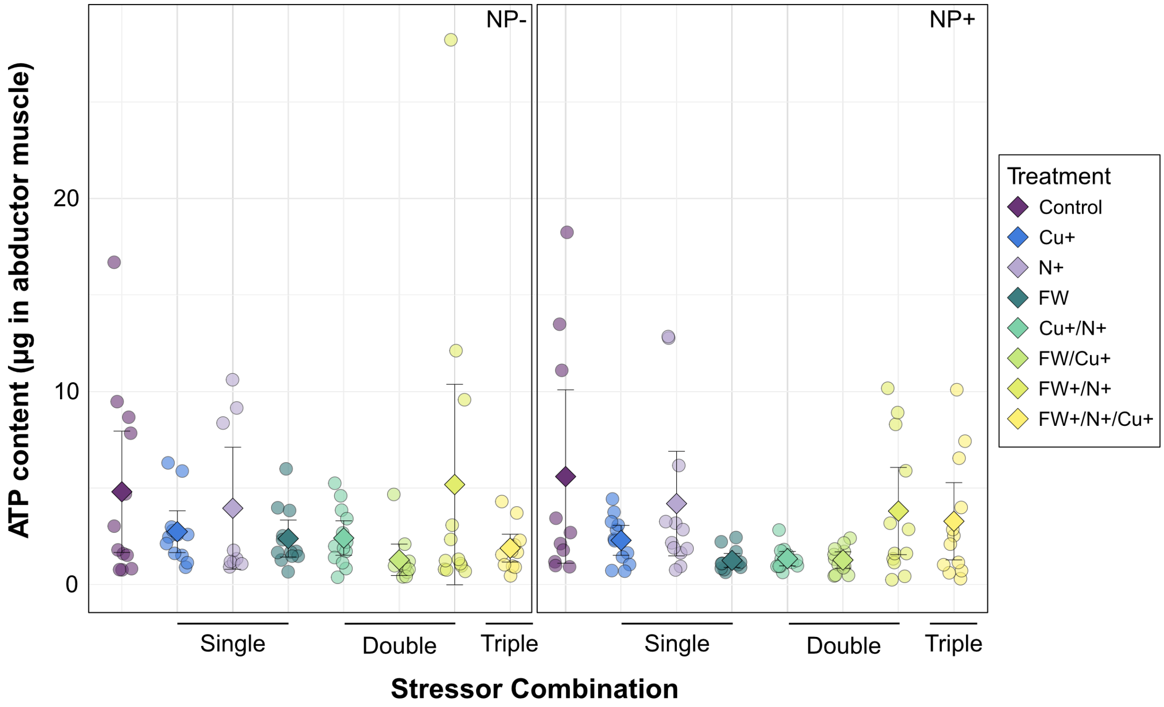


**Figure S8**. Mean ATP content (µg per abductor muscle) of *M. edulis* across all experimental conditions. Results are presented for individual and combined stressors, including copper contamination ($Cu^{+}$), nutrient enrichment ($N^{+}$), and freshwater input (FW), under conditions with and without nanoplastic pre-exposure ($NP^{+}$). Error bars represent 95% confidence intervals.

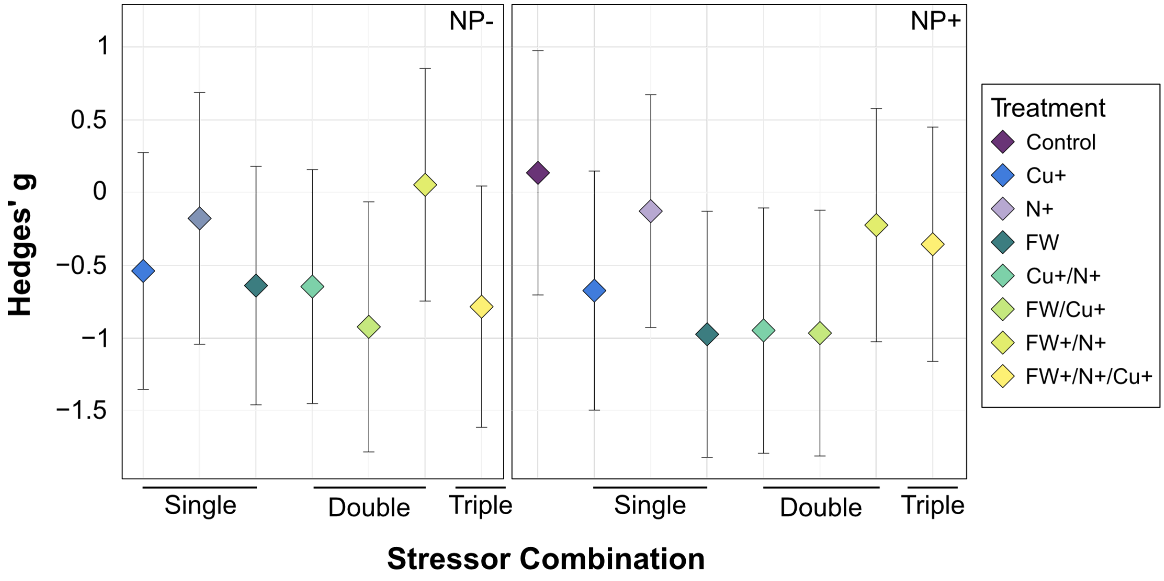


**Figure S9.** Standardized effect sizes (Hedges'g) comparing ATP content under individual and combined stressors, relative to control conditions (not shown on figure). Error bars correspond to 95% confidence intervals. Negative results show a decrease in survival rates compared to control conditions.
